# Rapid transcriptional response to physiological neuronal activity *in vivo* revealed by transcriptome sequencing

**DOI:** 10.1101/005876

**Authors:** Yarden Katz, Tarciso Velho, Vincent Butty, Christopher B. Burge, Carlos Lois

**Affiliations:** Dept. of Brain and Cognitive Sciences, MIT, Cambridge, MA; Dept. of Biology, MIT, Cambridge, MA; The Picower Institute for Learning and Memory, Cambridge, MA; Dept. of Biological Engineering, MIT, Cambridge, MA

## Abstract

Neuronal activity serves as a gateway between external stimulus from environment and the brain, often inducing gene expression changes. Alternative splicing (AS) is a widespread mechanism of increasing the number of transcripts produced from a single gene and has been shown to alter properties of neuronal genes, such as ion channels (Xie and Black 2001) and neurotransmitter receptors (Mu et al. 2003). Patterns of neural tissue-specific AS have been identified, often regulated by neuron-specific splicing factors that are essential for survival (Jensen et al. 2000; Li et al. 2007), demonstrating the importance of AS in neurons. *In vitro* studies of neuronal activity found AS changes in response to neuronal activity in addition to transcriptional ones, raising the question of whether such changes are recapitulated *in vivo* on behaviorally relevant timescales. We developed a paradigm for studying physiological neuronal activity through controlled stimulation of the olfactory bulb, and performed RNA-Seq transcriptome analysis of olfactory bulbs from odor-deprived and stimulated mice. We found that physiological stimulation induces large, rapid and reproducible changes in transcription *in vivo*, and that the activation of a core set of activity-regulated factors is recapitulated in an *in vitro* model of neuronal stimulation. However, physiological activity did not induce global changes in post-transcriptional mRNA processing, such as AS or alternative cleavage and polyadenylation. In contrast, analysis of RNA-Seq from *in vitro* stimulation models showed rapid activity-dependent changes in both transcription and mRNA processing. Our results provide the first genome-wide look at neuronal activity-dependent mRNA processing and suggest that rapid changes in AS might not be the dominant form of transcriptome alterations that take place during olfactory rodent behavior.

## 1 Introduction

Neuronal activity links environmental signals and the brain, often inducing activity-dependent gene expression changes that are essential for synaptic plasticity, wiring of the nervous system during development (Lin et al. 2008), and learning and memory (Flavell and Greenberg 2008). Several activity-dependent protein-coding genes have been identified, such as the protein-synthesis independent Immediate Early Genes (IEGs) (Flavell and Greenberg 2008), and the molecular mechanisms controlling their expression and activation of their targets have been studied intensively. Mouse knockout studies have shown that loss of IEGs like c-fos in specific brain regions results in neuronal cell death and interferes with learning and memory tasks, indicating that activity-dependent transcription is critical for brain function (Zhang et al. 2002). Furthermore, deletions and breakpoints in the activity-dependent transcriptional activator CREB-Binding Protein (CBP) cause Rubinstein-Taybi syndrome, underscoring the relevance of these factors for human genetic disease (Petrij et al. 1995; Flavell and Greenberg 2008).

Post-transcriptional mechanisms of controlling gene expression, such alternative splicing (AS) and alternative cleavage and polyadenylation (APA) have also been shown to be regulated by neuronal activity (Li et al. 2007; Flavell et al. 2008). AS provides a way of diversifying the proteome and has been shown to alter properties of neuronal genes, such as ion channels and neurotransmitter receptors. Patterns of neural tissue-specific of AS have been described, often regulated by neuron-specific expression of splicing factors (Li et. al., 2007). *In vitro* studies of neuronal activity documented post-transcriptional changes such as AS changes in response to neuronal activity in addition to transcriptional ones (Xie et. al. 2001, Ehlers et. al., Lee et. al. 2007) raising the question of whether such changes are recapitulated *in vivo* on behaviorally relevant timescales.

Activity-dependent gene expression is typically studied in culture by either KCl stimulation (Lin et al. 2008; Kim et al. 2010), long-term incubation with ion channel antagonists like bicuculline and tetrodotoxins (Ehlers 2003; Mu et al. 2003) or neurotransmitter receptor agonists (e.g. NMDA receptor agonists (Li et al. 2007)), potentially inducing neuronal depolarization at levels that are not representative of physiological activity in an intact brain. While there have been genome-wide studies of activity-induced gene expression *in vitro* (Lin et al. 2008; Flavell et al. 2008), the prevalence of activity-dependent post-transcriptional changes *in vivo* is unknown.

The olfactory bulb is an attractive model for studying neuronal activity-induced changes *in vivo*, since: (1) the synaptic pathway from olfactory sensory neurons in the nasal epithelium to the olfactory bulb is mapped and well-characterized, (2) the olfactory bulb is relatively homogeneous, easily dissectible brain region, made up of over 90% granule cells (Lledo et al. 2008), (3) olfaction is the primary sensory modality for rodents and is critical for survival, foraging and mating, and (4) the olfactory bulb is a highly conserved ancient brain region, similar in anatomy and function across vertebrates.

Here we use an olfactory stimulation/deprivation paradigm and high-throughput mRNA sequencing (RNA-Seq) to measure genome-wide activity-dependent changes in the olfactory bulb transcriptome *in vivo*. We found that physiological stimulation induces large, rapid and reproducible changes in transcription, and that the induction of a core set of activity-regulated factors is recapitulated in an *in vitro* model of neuronal stimulation. However, physiological neuronal activity did not result in global changes in post-transcriptional mRNA processing, such as AS or APA. In contrast, analysis of RNA-Seq from *in vitro* stimulation experiments shows rapid activity-dependent in both transcription and mRNA-processing.

## 2 Results

We developed a stimulation paradigm to induce neuronal activity in the olfactory bulb using odorants (Figure 1A). Control mice were kept in a sterile cage with a constant supply of odorless air (pumped through an activated carbon filter, described in Methods) for 12 hrs and then immediately sacrificed. To induce neuronal activity in the olfactory bulb, mice were exposed to a cocktail of odorants (listed in Supplementary Table 1) chosen to maximize activity across the olfactory bulb based on their empirical glumeruli activation profiles (Johnson and Leon 2007).

**Figure 1:**
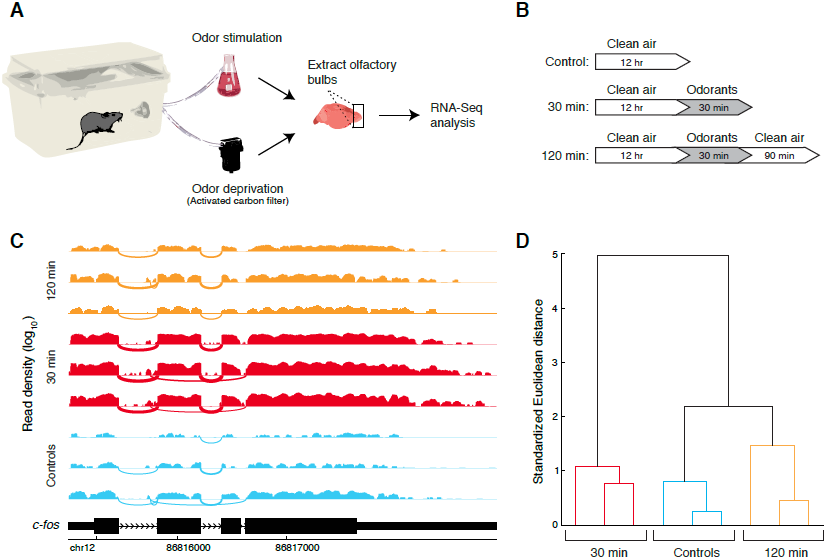
Olfactory stimulation paradigm followed by RNA-Seq. (A) Adult (8-11 weeks old) mice were kept in odor-free environment for 12 hrs using an activated-carbon air filter and then stimulated with a mixture of odorants (odor-deprived animals used as controls.) Immediately following stimulation or deprivation mice were sacrificed, and olfactory bulbs extracted from which RNA-Seq libraries were prepared. (B) Timelines of olfactory deprivation/stimulation experiments. Control animals were odor-deprived for 12 hours. Stimulated animals were odor-deprived for 12 hours, followed by either exposure to 30 minutes of odorant stimulation only (“30 min” time course) or 30 minutes of stimulation followed by 90 minutes of clean air (“120 min” time course). (C) RNA-Seq reads mapping to the c-fos immediate early gene from 9 mice (3 mice per condition) used in the experiment. Control condition replicates shown in blue, 30 min. condition in red and 120 min condition. Read densities correspond to number of reads overlapping a position (log_10_ units) and arcs correspond to exon-exon junctions whose width is proportional to number of junction reads. (D) Average-linkage hierarchical clustering on genes with fold change in expression greater than X between any pair of time points. Replicate mice cluster together, while the control gene expression profiles are globally more similar to 120 min mice than 30 min mice

**Table 1:**
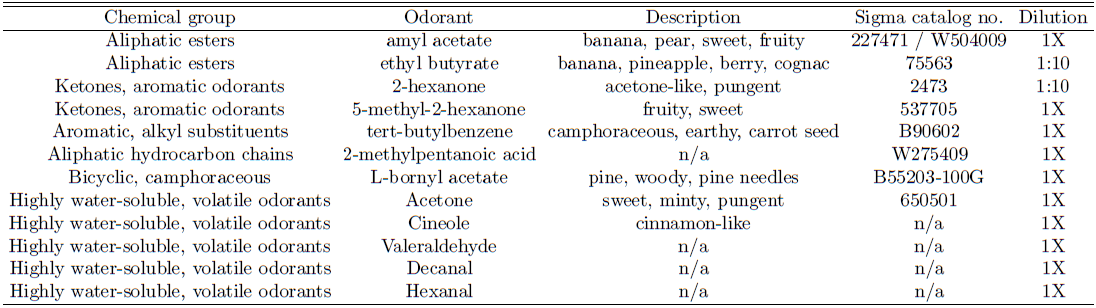
Odorant cocktail used to stimulate mouse olfactory bulb.

| Chemical group | Odorant | Description | Sigma catalog no. | Dilution |
| --- | --- | --- | --- | --- |
| Aliphatic esters | amyl acetate | banana, pear, sweet, fruity | 227471 / W504009 | 1X |
| Aliphatic esters | ethyl butyrate | banana, pineapple, berry, cognac | 75563 | 1:10 |
| Ketones, aromatic odorants | 2-hexanone | acetone-like, pungent | 2473 | 1:10 |
| Ketones, aromatic odorants | 5-methyl-2-hexanone | fruity, sweet | 537705 | 1X |
| Aromatic, alkyl substituents | tert-butylbenzene | camphoraceous, earthy, carrot seed | B90602 | 1X |
| Aliphatic hydrocarbon chains | 2-methylpentanoic acid | n/a | W275409 | 1X |
| Bicyclic, camphoraceous | L-bornyl acetate | pine, woody, pine needles | B55203-100G | 1X |
| Highly water-soluble, volatile odorants | Acetone | sweet, minty, pungent | 650501 | 1X |
| Highly water-soluble, volatile odorants | Cineole | cinnamon-like | n/a | 1X |
| Highly water-soluble, volatile odorants | Valeraldehyde | n/a | n/a | 1X |
| Highly water-soluble, volatile odorants | Decanal | n/a | n/a | 1X |
| Highly water-soluble, volatile odorants | Hexanal | n/a | n/a | 1X |

Two stimulation time courses were used to induce neuronal activity. The first wave of transcriptome changes was measured in a short time course, where animals were placed in the odorant-free cage for 12 hrs and then received 30 min of odorant stimulation. A second time course was used to measure effects downstream of the first wave and changes with slower kinetics, where mice received the same treatment as the 30 min animals but were given an additional 90 minutes of clean air (“120 min” time course.) (Figure 1B). Mice in the stimulated conditions exhibited highly exploratory behavior and movement upon induction of the odorants and throughout its duration, suggesting that animals were not desensitized to odorants during the experiment.

**Figure 2:**
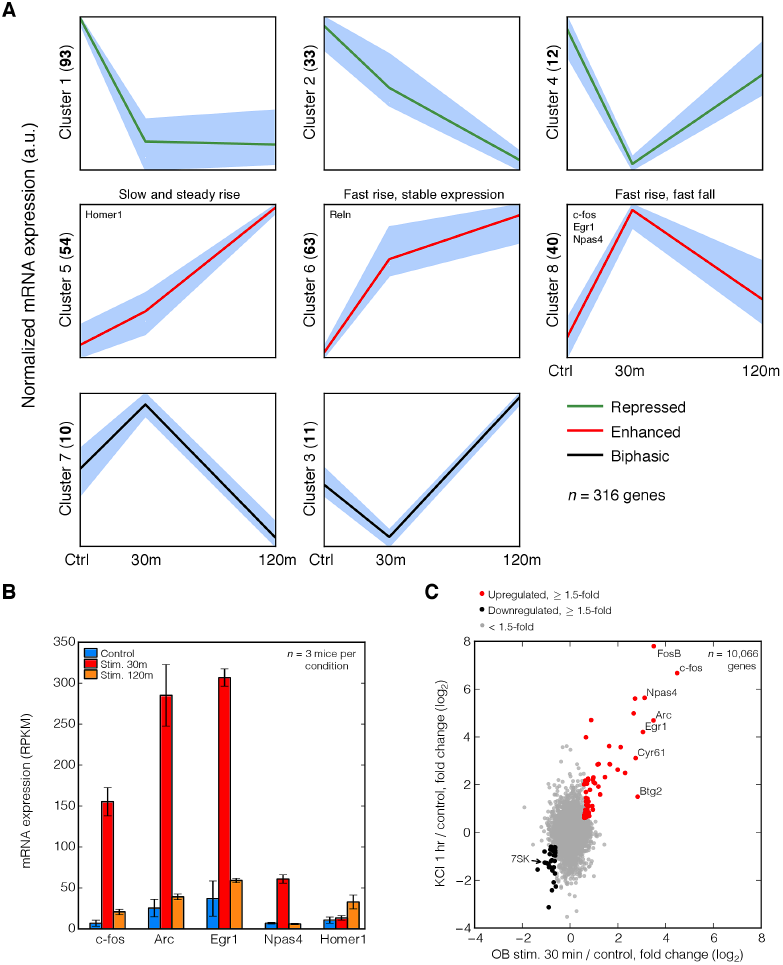
Transcriptional changes induced by activity. (A) Hierarchical clustering of gene expression profiles (average of replicates) across time points. Mean *z*-score normalized expression values are plotted along with shaded regions around standard deviation of scores. Mean expression of genes whose expression is enhanced by activity in both time points is shown in green (“Enhanced” genes), those whose expression is repressed in red (“Repressed” genes), and genes that change direction between the two time points in response to activity (relative to the control) in black (“Biphasic” genes.) Numbers in parentheses on y-axis of each cluster indicate the number of genes in that cluster. (B) Gene expression changes from olfactory bulb RNA-Seq for immediate early genes *c-fos*, *Arc*, *Egr1*, *Npas4* and *Homer1*. Expression levels measured in ‘reads per kilobase per million mapped reads’ (RPKM) in controls, 30 min stimulation and 120 min stimulation are plotted in blue, red and orange, respectively. (C) Scatter plot of fold changes (in log_2_) in expression between OB 30 min stimulation time point and OB control versus neuronal culture 1 hr KCl and culture controls. Grey points (plotted with alpha transparency) correspond to genes with fold changes smaller than 1.5 in either experiment, red to genes upregulated with fold changes ≥ 1.5-fold in both experiments, and black to genes downregulated with fold changes ≥ 1.5-fold in both experiments. Select commonly upregulated IEGs and downregulated 7SK noncoding RNA are labeled.

We prepared RNA-Seq libraries from whole olfactory bulbs (see Methods), obtaining a total of ∼320 M uniquely mapping reads from 9 mice. Reads were mapped to the genome and splice junctions (see Methods) and then used to compute gene expression profiles for control and stimulated mice. Immediate early genes (IEGs), such as c-fos, were rapidly induced in all 30 min stimulated mice, with upto ∼22 fold change in expression relative to controls, shown in (Figure 1C). Induction of the IEG Egr1 mRNA was confirmed by in situ hybridization (Supplementary Figure S1), corroborating RNA-Seq results, while induction of c-fos protein was confirmed by immunohistochemistry one hour following stimulation (data not shown.)

Hierarchical clustering of gene expression profiles for all mice revealed that replicate samples cluster together and exhibit high correlations in gene expression (Figure 1D). Furthermore, 120 min stimulated mice were more similar globally to the control mice than to 30 min stimulated mice, reflecting the fact that most activity-induced gene expression changes are transient and return to near-baseline levels by 120 min (Figure 1D).

To determine the overall kinetics of transcriptional changes, we performed hierarchical clustering analysis of gene expression profiles across the control and stimulation time points (Figure 2A). Several clusters of genes emerged that vary in the direction and shape of expression profiles. IEGs such as c-fos and Egr1 fit the expected “fast rise, fast fall” genes cluster, where expression is quickly upregulated but returns to baseline levels by 120 min (Figure 2A, Cluster 8). A distinct cluster of genes showing stable upregulation was also detected, and contained IEGs with slower kinetics like Homer1 (Figure 2A, Cluster 5).

**Figure 3:**
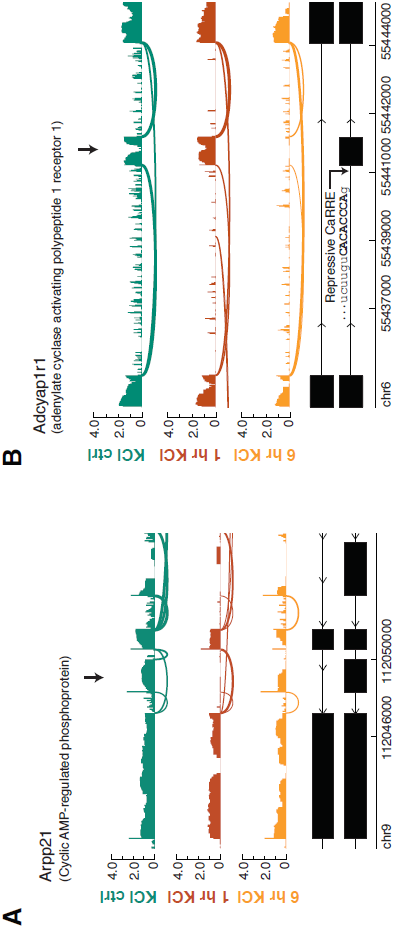
Activity-dependent splicing in neuronal culture stimulated with KCl. Effect of activity on mRNA-processing. (A) Rapid KCl-induced repression of alternative exon in *Arpp21* gene. Read densities (in base log_10_) along exons are plotted along with junctions as arcs. By 1 hr KCl, the alternative exon (marked by arrow) is nearly absent from transcripts. (B) Exon (marked by arrow) in *Adcyap1r1* is strongly skipped in 6 hr KCl condition, but not 1 hr KCl, relative to control. Upstream intron of repressed contains CaRRE (sequence shown in capital bold letters, marked by curved arrow) known to inhibit splicing in depolarization-dependent manner

Clusters were classified into three general categories: clusters where the net effect of activity is to upregulate expression (“Enhanced” genes, whose expression is increased in both stimulated conditions relative to controls), genes where the net effect of activity is to downregulate expression (“Repressed” genes, whose expression is decreased in both stimulated conditions), and genes where the effect of activity switches between the two stimulation conditions relative to control (“Biphasic” genes). This demonstrates heterogeneity in the effect of activity on transcription.

Known IEGs were induced reproducibly in all stimulated mice compared to controls (Figure 2B) and exhibited the expected kinetics. We also observed increased transcription of several activity-dependent factors implicated in neurogenesis/cell-cycle or enriched in newborn cells (Supplementary Figure S2), such as Reelin (Hack et al. 2002) and TenascinR (the latter known to be activity-dependent (Saghatelyan et al. 2004)), suggesting that whole olfactory bulb RNA-Seq provides sufficient sensitivity to detect changes in genes expressed in specific cell subtypes.

Recently, high-resolution RNA-Seq and ChIP-Seq was performed on mouse neurons stimulated *in vitro* look at activity-regulated RNAs and transcription factors controlling their expression (Kim et al. 2010). Cultured neurons were profiled following 1 and 6 hours of KCl treatment.

We applied the computational methodology used to analyze olfactory bulb RNA-Seq data to examine activity-regulated transcription and mRNA-processing in KCl-treated and control cultured neurons. Fold changes in expression between the first stimulated conditions and controls in the olfactory bulb and neuronal cultures were compared (using 30 min stimulation and OB control versus 1 hr KCl stimulation and KCl control, respectively, Figure 2C). As expected, the majority of genes show fold changes *<* 1.5-fold upon stimulation in both experiments. Strikingly, a group of genes were strongly upregulated—with fold changes upwards of 2-fold—in the stimulation conditions of both experiments (red points, Figure 2C). This group comprised the genes with the largest fold changes overall, though the magnitude of fold changes was consistently larger in neuronal culture. ChIP-Seq analysis showed that the majority of these commonly induced factors were bound by *Npas4*, consistent with the possibility that the maintenance of their expression is regulated in part by *Npas4* (data not shown.) We also noted a smaller number of genes that are downregulated with activity in both experiments.

Several novel predicted protein-coding genes were upregulated in both experiments (data not shown), while the noncoding small nuclear RNA 7SK, implicated in transcriptional control (Peterlin et al. 2012), was downregulated in both experiments. 7SK is traditionally thought to be transcribed by RNA Pol III, though recent evidence suggests it associates with RNA Pol II and has a histone signature of Pol II-transcribed loci (Barski, 2010). This suggests that regulatory noncoding RNAs also play a role in activity-dependent regulation in neurons.

We next asked whether activity induces changes in mRNA-processing. Several activity-dependent skipped exon events and their associated cis-regulatory elements have been identified (Li et al. 2007; Xie et al. 2005). It was also previously shown using exon arrays that long-term visual deprivation of mice followed by light stimulation induces changes in alternative cleavage and polyadenylation of a subset of transcripts in visual cortex (Flavell et al. 2008). Therefore, we focused on alternatively skipped exons and alternative cleavage and polyadenylation sites as representative instances of mRNA-processing events.

We found that activity did not induce global changes in olfactory bulb splicing patterns, and similar trends were observed for alternative cleavage and polyadenylation events (data not shown.) By contrast, the same analysis revealed strong and rapid activity-dependent splicing changes in neuronal cultures. The KCl-regulated exons we discovered corroborated findings of activity-dependent splicing in culture (Li et al. 2007; Xie et al. 2005). Certain exons showed strong and rapid switch-like changes in splicing within 1 hr of KCl treatment (Figure 3A,B), which is significantly quicker than the 24-48 hr incubations with depolarizing agents used for previous *in vitro* measurements of activity-dependent splicing (Lee et. al., 2007). Some exons were also strongly regulated by activity in the 6 hr KCl time point. The *Adyacap1r1* gene contains a calcium responsive RNA element (CaRRE) in the intron upstream of Exon 14 (Figure 3B) that causes repression of the exon upon depolarization in CaMK IV-dependent manner (Li et al. 2007). We found that this exon is strongly skipped with activity and is near absent in the 6 hr KCl condition (Figure 3C). Similarly, *Arghgap21* contains an exon that decreases ∼3-fold in inclusion levels by 6 hours of KCl treatment (Supplementary Figure S2).

## 3 Discussion

Our study provides an *in vivo* genome-wide look into physiological activity-dependent mRNA processing in the mouse olfactory bulb transcriptome. Our results suggest that rapid transcriptional regulation, but not mRNA-processing, predominates the olfactory bulb transcriptome response to neuronal activity. This has implications for the study of gene expression changes induced by olfaction-based behavioral experiments in rodents.

A systematic comparison of our *in vivo* stimulation of olfactory bulbs and a commonly used *in vitro* model of neuronal stimulation suggests that a core set of activity-regulated factors is shared across diverse neuronal cell types, but that mRNA-processing changes are not. Recent work shows that not all brain regions employ a high degree of alternative splicing. For example, recent study of cerebellum (Pal et. al., 2011) showed that throughout development of the brain region, alternative promoter usage is a much more commonly used form of generating gene product diversity, compared with alternative splicing. Both the cerebellum and the olfactory bulb are ancient and highly conserved brain regions in vertebrates, leaving open the possibility that evolutionarily newer regions (like cortex) do employ alternative splicing and mRNA-processing more widely.

We cannot exclude the possibility that some of the discordance observed between our *in vivo* model and *in vitro* stimulation models are due to the difference in developmental stages of the cells (neuronal cultures originating from embryo, our tissues from an adult animal.) In principle, there could be developmental stage-specific activity-dependent instances of mRNA-processing. Future studies using transcriptome sequencing in of activated neurons in distinct developmental stages and brain regions could resolve these issues.

## 4 Methods

### 4.1 Olfactory stimulation and deprivation setup

C57BL6 mice (Charles River laboratory) ages 8-11 weeks were placed in an autoclaved ethanol-sterilized cage and placed in a clean fume hood for 12 hours overnight, with a steady supply air filtered by a compressed air activated-carbon filter (Hankison, Inc.). Filter was hooked into laboratory compressed air nozzle mediated by an air pressure gauge. A timer-controlled light source programmed to the animal housing facility’s light/dark cycle was placed above the cage during experiments. All animals were sacrificed between 10 am and 12 pm.

### 4.2 Odorants preparation

A set of 12 odorants (listed in Supplementary Table 1) was selected based on each odorant’s activation profile of specific glomeruli or subregions of the olfactory bulb as described in (Johnson and Leon 2007) and related studies. Odorant mixtures were comprised both glomerulus-specific and broad activators of the olfactory bulb (such as valeric acid and methyl valerate), with many of the odorants having food-associated scents. Odorants were diluted in Light Mineral Oil (Sigma) and pipetted onto damp pieces of filter inside a capped Erlenmeyer flask, which was plugged into the air supply of the mouse cage during stimulation and pumped in at 3 psi.

### 4.3 Tissue preparation and extraction of RNA

From each animal, one hemisphere’s olfactory bulb (OB) was used for RNA extraction and RNA-Seq library preparation and the remaining bulb taken for RNA in situ hybridization and immunohistochemistry to confirm induction of immediate early genes *Egr1* and *c-fos*. OBs for RNA-Seq were dissociated in Trizol (Invitrogen) and RNA extracted using standard Trizol extraction, followed by a run on a BioAnalyzer to profile RNA quality and obtain RNA Integrity Number (RIN). The remaining intact OBs (attached to forebrain) were prepared for sectioning by immersion in TissueTek inside a plastic well, and then flash frozen in crushed dry ice and isopropyl alcohol. Cryostat coronal sections of frozen OBs were prepared for in situ hybridization.

### 4.4 Sample preparation for RNA-Seq

RNA-Seq libraries were prepared based on protocols described elsewhere (Mortazavi et al. 2008). Briefly, two rounds of polyA+ RNA selection were performed using poly(T) beads. Remaining polyA+ RNA was fragmented and prepared for Illumina sequencing: paired-end adaptors were ligated onto cDNAs primed with random hexamers and products were size-selected by agarose gel electrophoresis. Bands ranging from ∼250-300 bp were cut out using GeneCatcher gel excision tips and purified. Samples were PCR amplified for 13 cycles prior to sequencing.

### 4.5 Mapping and computational analysis of RNA-Seq data

Reads were mapped to mouse genome (mm9) and a set of splice junctions (Wang et al. 2008) using bowtie (Langmead et al. 2009). Only uniquely mapped reads were used for downstream analyses. Quantitation of exon abundances was performed using MISO (Katz et al. 2010). Gene expression levels quantitated by computing RPKM values (Mortazavi et al. 2008) using only reads from constitutive exons.

SOLiD mRNA-Seq sequencing data from (Kim et. al., 2010) was obtained from GEO (Accession code: GSE21161) and mapped to the same mouse (mm9) indexed genome plus splice junctions, built using the bowtie color index (-C) option.

Expression profiles of genes (in RPKM units) showing 1.5-fold change or greater between any pair of olfactory bulbs were used to perform hierarchical clustering using the Euclidean distance metric in Figure 1D. These expression profiles were z-score normalized to perform hierarchical clustering analysis of each gene’s expression profile over the three olfactory bulb conditions (Figure 2A). All genes expression analyses and clustering were performed using custom-built scripts written in the Python programming language, using scipy/numpy packages.

### 4.6 Data availability

mRNA-Seq data from this study are available at GEO (Accession code: GSE58843).

## 5 Acknowledgements

We thank Douglas Black and Yingxi Lin for helpful discussion.

## 6 Author contributions

YK, TV, and CL conceived the project and designed experiments. YK and TV performed mouse experiments and prepared RNA-seq libraries. YK performed computational analyses. VB performed computational analyses of RNA editing and contributed expert advice. CB contributed to computational analyses. mRNA-Seq library preparation and sequencing took place 2009-2010.

---

**Supplementary Figure S1:**
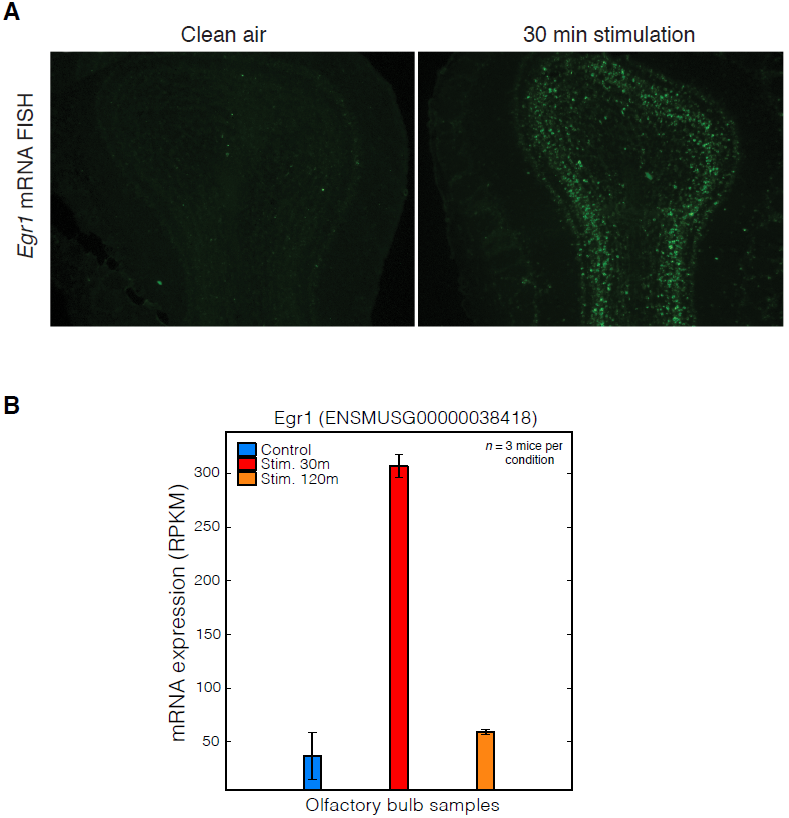
Activation of immediate early genes validated by in situ hybridization. (A) mRNA fluorescent in situ hybridization for *Egr1* in coronal olfactory bulb sections from control (“clean air”) and 30 min. stimulated mice. (B) RNA-Seq estimates of gene expression (in RPKM) for *Egr1* in controls and stimulated mice.

**Supplementary Figure S2:**
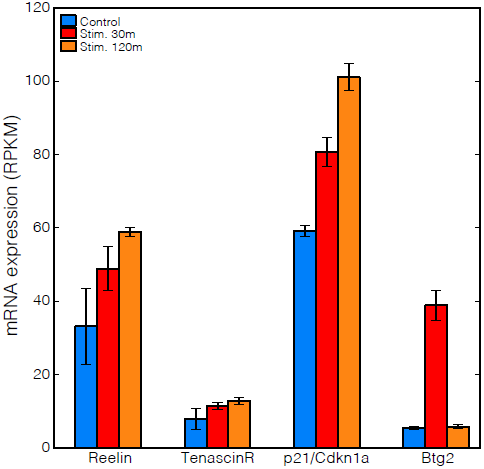
Activation of adult neurogenesis and neuronal migration genes. RNA-Seq expression levels for genes either associated with adult neurogenesis or enriched in newborn/migrating cells (*Reelin, TenascinR, p21/Cdkn1a, Btg2*) that show activity-dependent regulation in olfactory bulb samples.

## References

1. Ehlers, MD. Activity level controls postsynaptic composition and signaling via the ubiquitin-proteasome system. Nature neuroscience 6 (2003), pp. 231–242 (cit. on p. 3).

2. Flavell, SW and ME Greenberg. Signaling mechanisms linking neuronal activity to gene expression and plasticity of the nervous system. Annual review of neuroscience 31 (2008), pp. 563–590 (cit. on p. 2).

3. Flavell, SW, TK Kim, JM Gray, DA Harmin, M Hemberg, EJ Hong, E Markenscoff-Papadimitriou, DM Bear, and ME Greenberg. Genome-wide analysis of MEF2 transcriptional program reveals synaptic target genes and neuronal activity-dependent polyadenylation site selection. Neuron 60 (2008), pp. 1022–1038 (cit. on pp. 2, 3, 6).

4. Hack, I, M Bancila, K Loulier, P Carroll, and H Cremer. Reelin is a detachment signal in tangential chain-migration during postnatal neurogenesis. Nature neuroscience 5 (2002), pp. 939–945 (cit. on p. 5).

5. Jensen, KB, BK Dredge, G Stefani, R Zhong, RJ Buckanovich, HJ Okano, YY Yang, and RB Darnell. Nova-1 regulates neuron-specific alternative splicing and is essential for neuronal viability. Neuron 25 (2000), pp. 359–371 (cit. on p. 1).

6. Johnson, BA and M Leon. Chemotopic odorant coding in a mammalian olfactory system. The Journal of comparative neurology 503 (2007), pp. 1–34 (cit. on p. 3).

7. Katz, Y, ET Wang, EM Airoldi, and CB Burge. Analysis and design of RNA sequencing experiments for identifying isoform regulation. Nature Methods 7 (2010), pp. 1009–15 (cit. on p. 8).

8. Kim, TK, M Hemberg, JM Gray, AM Costa, DM Bear, J Wu, DA Harmin, M Laptewicz, K Barbara-Haley, S Kuersten, E Markenscoff-Papadimitriou, D Kuhl, H Bito, PF Worley, G Kreiman, and ME Greenberg. Widespread transcription at neuronal activity-regulated enhancers. Nature 465 (2010), pp. 182–187 (cit. on pp. 3, 5).

9. Langmead, B, C Trapnell, M Pop, and SL Salzberg. Ultrafast and memory-efficient alignment of short DNA sequences to the human genome. Genome Biol 10 (2009), R25 (cit. on p. 8).

10. Li, Q, JA Lee, and DL Black. Neuronal regulation of alternative pre-mRNA splicing. Nature Reviews Neuroscience 8 (2007), pp. 819–831 (cit. on pp. 1–3, 6).

11. Lin, Y, BL Bloodgood, JL Hauser, AD Lapan, AC Koon, TK Kim, LS Hu, AN Malik, and ME Greenberg. Activity-dependent regulation of inhibitory synapse development by Npas4. Nature 455 (2008), pp. 1198–1204 (cit. on pp. 2, 3).

12. Lledo, PM, FT Merkle, and A Alvarez-Buylla. Origin and function of olfactory bulb interneuron diversity. Trends in neurosciences 31 (2008), pp. 392–400 (cit. on p. 3).

13. Mortazavi, A, BA Williams, K McCue, L Schaeffer, and B Wold. Mapping and quantifying mammalian transcriptomes by RNA-Seq. Nature methods 5 (2008), pp. 621–628 (cit. on p. 8).

14. Mu, Y, T Otsuka, AC Horton, DB Scott, and MD Ehlers. Activity-dependent mRNA splicing controls ER export and synaptic delivery of NMDA receptors. Neuron 40 (2003), pp. 581–594 (cit. on pp. 1, 3).

15. Peterlin, BM, JE Brogie, and DH Price. 7SK snRNA: a noncoding RNA that plays a major role in regulating eukaryotic transcription. Wiley interdisciplinary reviews. RNA 3 (2012), pp. 92–103 (cit. on p. 5).

16. Petrij, F, RH Giles, HG Dauwerse, JJ Saris, RC Hennekam, M Masuno, N Tommerup, GJ van Ommen, RH Goodman, and DJ Peters. Rubinstein-Taybi syndrome caused by mutations in the transcriptional co-activator CBP. Nature 376 (1995), pp. 348–351 (cit. on p. 2).

17. Saghatelyan, A, A de Chevigny, M Schachner, and PM Lledo. Tenascin-R mediates activity-dependent recruitment of neuroblasts in the adult mouse forebrain. Nature neuroscience 7 (2004), pp. 347–356 (cit. on p. 5).

18. Wang, ET, R Sandberg, S Luo, I Khrebtukova, L Zhang, C Mayr, SF Kingsmore, GP Schroth, and CB Burge. Alternative isoform regulation in human tissue transcriptomes. Nature 456.7221 (2008), pp. 470–6 (cit. on p. 8).

19. Xie, J and DL Black. A CaMK IV responsive RNA element mediates depolarization-induced alternative splicing of ion channels. Nature 410 (2001), pp. 936–939 (cit. on p. 1).

20. Xie, J, C Jan, P Stoilov, J Park, and DL Black. A consensus CaMK IV-responsive RNA sequence mediates regulation of alternative exons in neurons. RNA (New York, N.Y.) 11 (2005), pp. 1825–1834 (cit. on p. 6).

21. Zhang, J, D Zhang, JS McQuade, M Behbehani, JZ Tsien, and M Xu. c-fos regulates neuronal excitability and survival. Nature genetics 30 (2002), pp. 416–420 (cit. on p. 2).

